# Enriched environmental conditions modify the gut microbiome composition and fecal markers of inflammation in Parkinson’s disease

**DOI:** 10.1101/614834

**Authors:** Yogesh Singh, Mohamed El Hadidi, Jakob Matthes, Zinah Wassouf, Julia M. Schulze-Hentrich, Ursula Kohlhofer, Leticia Quintanilla-Martinez de Fend, Daniel Huson, Olaf Riess, Nicolas Casadei

**Affiliations:** Institute of Medical Genetics and Applied Genomics, Calwerstrasse 7, 72076, Tuebingen, University of Tuebingen, Germany; Algorithms in Bioinformatics, Faculty of Computer Science, Sand 14, 72076, Tuebingen, University of Tuebingen, Germany; Bioinformatics, Center for Informatics Science, Nile University, Giza, Egypt; Institute of Pathology, University Hospital & Comprehensive Cancer Center, Liebermeisterstrasse 8 University of Tuebingen, Tuebingen, Germany; DFG NGS Competence Center, University of Tübingen, Tübingen, Germany

**Keywords:** alpha-synuclein, Parkinson’s disease, enriched environment, microbiome, inflammation

## Abstract

**Background:** Recent findings suggest an implication of the gut microbiome in Parkinson’s disease patients. Parkinson’s disease onset and progression has also been linked with various environmental factors such as physical activity, exposure to pesticides, head injury, nicotine, and dietary factors.

**Objectives:** In this study, we used a transgenic mouse model overexpressing the complete human SNCA genes modeling familial and sporadic forms of Parkinson’s disease to study whether environmental conditions such as standard *versus* enriched environment changes the gut microbiome and influences disease progression.

**Methods:** We performed 16S rRNA DNA sequencing on fecal samples for microbiome analysis and studied fecal inflammatory calprotectin from the colon of control and transgenic mice kept under standard environment and enriched environment conditions.

**Results:** The overall composition of the gut microbiota was not changed in transgenic mice compared with wild-type in enriched environment, however, individual gut bacteria at genus level such as Lactobacillus sp. were significantly changed in transgenic mice. Furthermore, enriched environment significantly reduced colon fecal inflammatory calprotectin protein in wild-type and transgenic enriched environment conditions compared to standard environment.

**Conclusion:** Our data suggest that enriched social environment has a positive effect on the induction of SNCA mediated inflammation in the intestine by changing anti-inflammatory gut bacteria.

## Introduction

Chronic inflammation is a key process in the progression of Parkinson’s disease (PD)^1^. Besides the main pathological features of Lewy bodies (LBs) and neurites predominantly composed of α-Synuclein (α-Syn) protein^2,3^ in the brain and gut of PD patients, neuroinflammatory markers such as reactive microglial expression of HLA-DR, CXCL2 and S100b in substantia nigra (SN), IgG in LBs, proinflammatory cytokines (IL1β and TNF-α) in SN and striatum are additional neuropathological characteristics of PD^4^. Interestingly, although multiple studies showed a progression of the pathology (also termed either α-Synucleinopathy or Synucleinopathy) through the brain^5,6^, other studies also proposed a possible spread of the disease from neurons to neurons *via* seed of α-Syn in the peripheral nervous system^7,8^. Accordingly, injection of α-Syn in the brain^9^, in the gut^10^ or in muscles^11^ pointed out a retrograde transport of protein seeds from the peripheral nervous system to the central nervous system. Thus, in addition to the genetic factors, external or unknown internal factors inducing α-Syn aggregation or expression such as environmental conditions, diet or lifestyle, bacterial metabolites and self-antigens may also trigger or enhance PD^12–20^.

Environmental factors appeared to improve and worsen the pathology of PD^15,20–22^. Several chemicals or pesticides such as MPTP^23^, paraquat^24,25^, rotenone^26^, viral infections^27^ etc. exposure lead to neurodegenerative effects whilst enriched environmental (EE) conditions based on studies of animal models suggested that enrichment can promote neuronal activation, signaling and plasticity in various brain regions. Therefore, EE conditions could possibly exert neuroprotective effects on neurons and delay pathogenesis in PD^12,28,29^.

The gut harbors a dynamic environment and is exposed to environmental factors such as diet, antibiotics, and pathogens and follows on constant interaction with microbial communities^30–32^. In addition, the gut microbiota is also shaped throughout life by host-related factors such as host genotype as well as with ageing^33^. Disturbances within gut microbiota have been reported to influence host susceptibility to pathogens and pathological conditions such as gastrointestinal inflammatory diseases and obesity^34–37^. Gastrointestinal dysfunctions are frequently reported by PD patients^38^ and in some clinical studies are preceding the onset of motor symptoms^39,40^. In this context, the presence of Synucleinopathy in the autonomic nervous system of PD patients^41^, accumulation of α-Syn in the bowel of PD patients^42^ as well as a modified intestinal microbiota correlated to motor symptoms suggest a potential origin of Synucleinopathy in the gastrointestinal tract (GIT)^43^. Several recent patients from different countries (USA, Japan, China, Germany, and Netherlands) and mouse studies suggest a potential role of gut microbiome in PD pathogenesis^43–51^ as gut dysbiosis may lead to an inflammatory environment potentially initiating synucleinopathy^52–54^.

In this study, we aim to investigate if the expression of α-Syn in the gut of a humanized mouse model of PD can be influenced by EE conditions and explore how EE conditions interact with the gut microbiome to affect the inflammation in the gut.

## Methods

### Animal experiments and EE conditions

C57BL/6N mice were obtained from Charles River and were housed in colony cages under a 12 hour light-dark cycle with free access to food and water. BAC-hSNCA Transgenic (SNCA-TG) mice and EE conditions have been described earlier^55^. To assess the impact of α-Syn and enriched environmental conditions on microbiome composition changes, 12-month-old female mice were used.

### Euthanasia and sample collection

To minimize the influence of preparation on the gut microbiome, mice were euthanized via cervical dislocation during the light phase. GIT was then prepared rapidly on ice, content of the gut as well as the rest of the tissue rinsed in PBS (10 mM phosphate, 150 mM NaCl, pH 7.4) were collected in cryotubes and snap frozen by liquid N_2_ and stored at −80°C until use.

### DNA isolation, preparation and sequencing

DNA was isolated from the fecal and cecal content. Briefly, 10 to 20 mg of digesta was mixed with 180 μl of 20 mg/ml lysozyme (L6876; Sigma) in 20 mM TrisHCl at pH 8.0, 2 mM EDTA and 1.2% of triton-X. The suspended pellet was incubated under agitation for 30 min at 37 °C and then digested at 56°C for 30 min with 20 μl of 50 mg/ml proteinase K (315836; Roche) in 200 μl of buffer AL. Digestion was inactivated at 95°C for 15 min. After centrifugation, 200 μl of ethanol was added to the samples prior to binding to QIamp spin column (QIAamp DNA Blood mini kit; Qiagen) and samples were centrifuged at 8000 rpm for 1 minute. Samples were then washed in successively 500 μl of buffer AW1 and AW2. DNA was then eluted in 200 μl of buffer AE and stored at −20°C.

For 16S rRNA amplification, 12.5 ng of DNA was amplified using 0.2 μM of both forward primer (TCGTCGGCAGCGTCAGATGTGTATAAGAGACAGCCTACGGGNGGCWGCAG, Metabion) and reverse primer (GTCTCGTGGGCTCGGAGATGTGTATAAGAGACAGGACTACHVGGGTATCTAATCC, Metabion) and KAPA HiFi HotStart Ready Mix (KK2601; KAPABiosystems). PCR was performed using a first denaturation of 95°C for 3 minutes (min), followed by 25 cycles of amplification at 95°C for 30 s, 55°C for 30 s and 72°C for 30s, final elongation at 72°C for 5 min and the amplified DNA was stored at 4°C. Electrophoresis of samples was used to verify amplicon specificity.

Samples were then purified (Agencourt AMPure XP, Beckman Coulter) and PCR amplicons were indexed using Nextera XT index and KAPA HiFi HotStart ReadyMix. PCR was performed using a first denaturation of 95°C for 3 min, followed by 8 cycles of amplification at 95°C for 30 s, 55°C for 30 s and 72°C for 30, final elongation at 72°C for 5 min. Samples purified were then validated using BioAnalyzer (Bioanalyzer DNA 1000, Agilent) and 4 nM of each library pooled using unique indices before sequencing on a MiSeq (Illumina) and paired 300-bp reads.

### Sequence Analysis and Statistics

Available sequence data has been trimmed and filtered using SeqPurge^56^. Trimming parameters demanded a minimum quality at 3’ end of q=35 (parameter qcut=35). Processed sequence data has been aligned using MALT (version 0.3.8; https://ab.inf.uni-tuebingen.de/software/malt) against the 16S database SILVA SSU Ref Nr 99 (https://www.arb-silva.de/documentation/release-128/) and classified using NCBI taxonomy. Alignment has been performed using semi-global alignment and a minimum sequence identity of 90% (parameter minPercentIdentity=90). Further analysis and visualization was performed using MEGAN (MEtaGenome Analyzer) version 6.0^57^. In Figure 4C, data were normalized with SE and EE conditions and shown as percentage abundance. Alpha (statistical) was set a priori at 0.05 for all tests of significance.

### Preparation of tissue for histological analysis

Gut samples prepared on ice were fixed for 24 h in 4% paraformaldehyde (PFA), stored at 4°C in 0.4% PFA for a maximum of 4 weeks prior embedding in paraffin. Brain samples were prepared from mice deeply anesthetized by CO2 inhalation and transcardially perfused with PBS and cold 4% PFA diluted in PBS. Brains were removed carefully from the skull and post-fixed 24 h in 4% PFA. Fixed samples were then alcohol-dehydrated and embedded in paraffin. Samples were embedded in paraffin blocks using a tissue embedding station and stored at room temperature until use. Paraffin blocks containing colon and other GIT tissues were cut into 7 μm thick sections using a microtome. Section were placed in 45°C water bath for flattering, collected on a glass slide, dried in an incubator at 50°C for 1 h and stored at room temperature.

### Immunohistochemistry

Immunohistochemistry was performed on colon sections as described earlier on brain sections^58^ using additional antibodies (α-Syn: ab27766, Abcam or L001, Enzo; NeuN: MAB377, Merck; MAP2: ab70218 Abcam; TH: 657012, Merck; GFAP: Z0334, Dako).

### Tissue lysate

Gut tissue were weighted frozen and lysed with 10 volumes of RIPA buffer (50 mM Tris, 150 mM NaCl, 1.0% NP-40, 0.5% sodium deoxycholate, 0.1% SDS, pH 8.0) supplemented with protease inhibitor (Complete; Roche Diagnostics). Brain tissues were homogenized 30 sec using a disperser (T10 ULTRA-TURRAX; VWR) on ice. After the homogenization, samples were incubated for 30 min at 4°C and spun for 20 min at 12 000 g. Proteins lysate supernatants were supplemented with 10% glycerol before long storage at −80°C. Protein concentration was determined using BCA method (23225; Life Technology).

### Western blotting

Samples were prepared by diluting protein lysates in PAGE buffer (0.2 M glycine, 25 mM Tris, 1% SDS), followed by a denaturation at 95°C for 10 min in loading buffer (80 mM Tris, 2% SDS, 5% 2-mercaptoethanol, 10% glycerol, 0.005% bromophenol blue, pH 6.8) and a short centrifugation 30 sec at 400 g. Proteins were separated by electrophoresis using 12% SDS-PAGE gel. Gels containing proteins were washed for 5 minutes in transfer buffer (0.2 M glycine, 25 mM Tris, 10–20% methanol) and transferred to membranes equilibrated in transfer buffer. Transfer was performed for 90 min at 80 V at 4°C on nitrocellulose membranes (88018, Life Technology). Immunoblot were washed 5 min in TBS buffer and blocked using 5% non-fat milk (Slim Fast) in TBS. Membranes were then washed twice 5 min in TBST and incubated with the primary antibody over night at 4°C (human and mouse a-syn: 610786 BD Biosciences; human α-syn: 804-258-1001, Enzo Life Science; beta-actin: A4700, Sigma). After incubation with the first antibody, membranes were washed four times (5 min each) with TBST. Membranes were then incubated for 75 min with the secondary antibody coupled to horseradish peroxidase (GE Healthcare). After four washing steps with TBST (5 min each), bands were visualized using the enhanced chemiluminescence method (ECL+; GE Healthcare). Light signal was detected using LI-COR Odyssey and were quantified using Odyssey software.

### Calprotectin ELISA

To measure the calprotectin/MRP 8/14 from the fecal samples, S100A8/S100A9 ELISA kit (#K6936, Immundiagnostik AG) was used according to the manufacture’s guidelines. First fecal samples were measured (weight between 20 mg and 50 mg) and dissolved in 500 ul of extraction buffer supplied by the kit, mixed by vortexing and then centrifuged for 10 min at 3000 g. Supernatant was taken and transferred to a new microcentrifuge tube and 100 ul of sample was used for measuring the protein. The data was analyzed using 4 parameters algorithm and the concentration of calprotectin was normalized with feces weight and data are presented in ng/ml/g.

### Statistical analysis

MEGAN-CE (version 6.10.6, built 20 Dec 2017) was used for data acquisition and analysis. GraphPad and Inkscape were used for the final figure preparation. One way ANOVA and student’s t-test was used for statistical analysis using GraphPad. The p value (≤0.05) considered significant.

## Results

### Human α-Syn expression in the enteric nervous system (ENS) of the gut in SNCA-TG mice

To study if the expression of α-Syn may induce changes of the gastrointestinal microbiome, we used a SNCA-TG mouse model of Parkinson’s disease overexpressing the complete human *SNCA* gene including its promoter, introns, exons and UTRs^59^, as reported previously^55^. This model is overexpressing the human protein with a spatial distribution in the brain similar to the endogenous human and mouse expression^55^. Using immunohistochemistry (IHC), we investigated the expression of the transgenic α-Syn in the gastrointestinal tract of the SNCA-TG mice and observed the presence of the human α-Syn protein in the colon and the entire gut including stomach, ileum, cecum, and rectum, respectively (Fig. 1A and Suppl. Fig. 1). Staining was observed in the muscular layer and more particularly in the longitudinal muscle. Interestingly, we also observed a specific staining in the mucosal layer of the colon. Double immunofluorescence stainings were performed using antibodies against human α-Syn and neuronal markers such as map2, neurofilament, and NeuN. Staining identified double-labeled neurons positive for human α-Syn protein and is confirming a strong expression of the transgenic protein in the ENS (Fig. 1B). A strong expression of human α-Syn was also confirmed using Western blots (Figure 1C). The level of total α-Syn observed in SNCA-TG mice was significantly increased in SNCA-TG mice compared to WT in both, the caecum and the colon (Fig. 1C). Thus, these results suggested that our SNCA-TG animal model is suitable to study the role of SNCA overexpression in the gut and to study host-pathogen interactions for the impact of EE conditions on the accumulation of α-Syn.

**Fig. 1.**
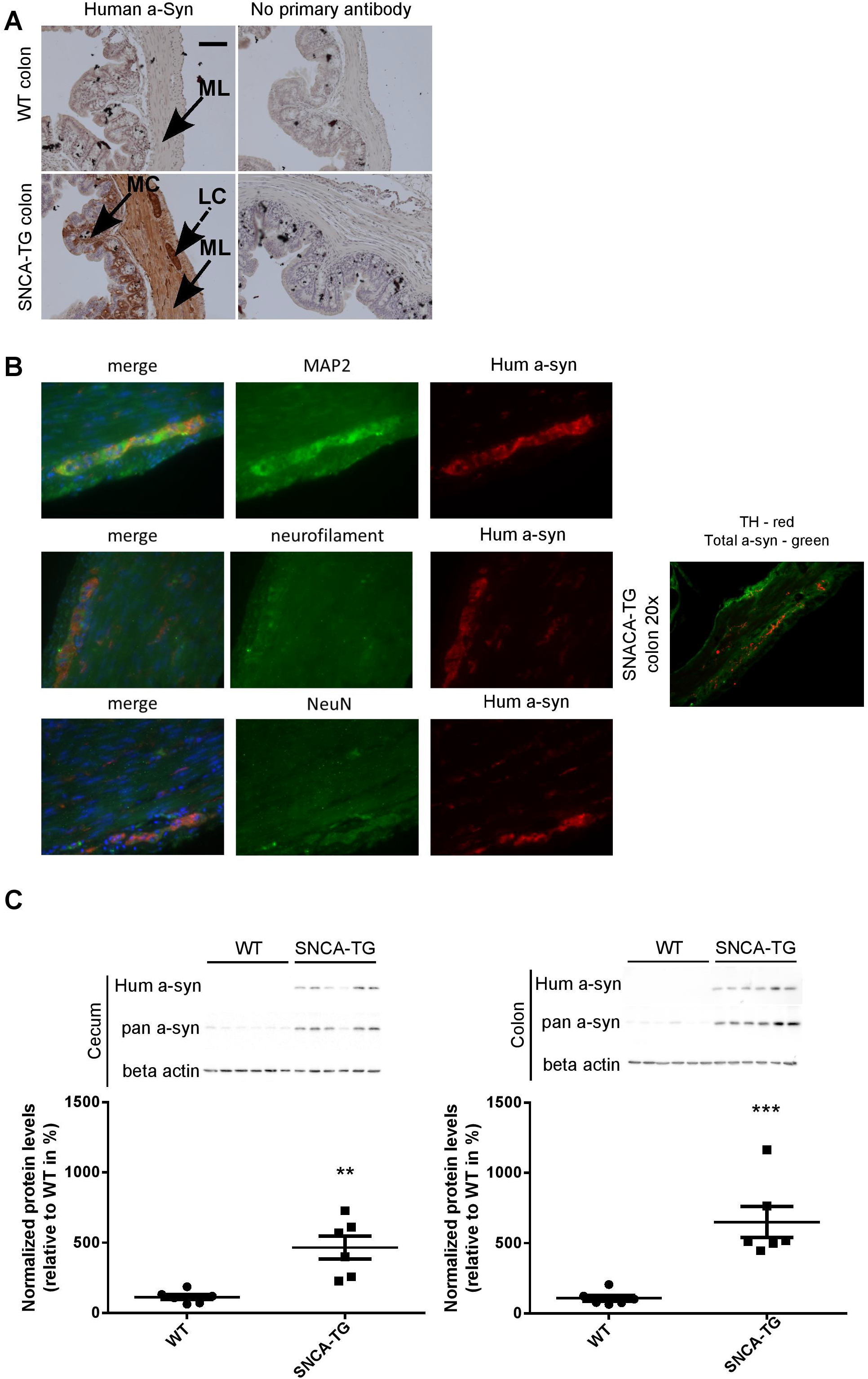
Human α-Syn expression in the ENS (colon) of SNCA-TG mice. Expression of the transgenic human α-Syn protein was investigated by IHC and Immunoblotting (A). Transgenic human α-Syn protein was detected in the gastrointestinal tract of transgenic mice using IHC in muscular layer, longitudinal muscle cells, and mucosal layer of the colon. (A). (B) Double immunofluorescence staining suggests the presence of transgenic α-Syn in enteric neurons labeled with the synaptic markers MAP2, the neuronal markers neurofilament (NeuN), and the dopaminergic marker TH. (C) Expression of human α-Syn in the gut was confirmed using Western blotting from 12month-old mice (n=6). Unpaired two-tailed t test was performed the significance measurements. P value was considered significance when it was less than or equal to 0.05 (p ≤ 0.05*, ** p≤0.01, *** p≤0.001).

### Changes in the gut microbiome composition in SNCA-TG mice under SE conditions

To compare gut microbial communities diversity, we isolated DNA from 12 month-old WT and SNCA-TG caecal and colon mouse digesta, performed PCR of the 16S ribosomal RNA (16S rRNA) and sequenced the amplicons using next generation sequencing^60,61^. Due to the elevated expression of SNCA and the involvement of constipation in the mice overexpressing SNCA^62,63^, we focused on the cecum and colon microbiota and sequenced both groups with a comparable depth of ~100,000 clusters per sample (Suppl. Fig. 2A and Suppl. Fig. 3A). We then evaluated the population diversity of the microbial community (alpha-diversity) using Shannon-Weaver index (SWI) and found no significant differences between the WT and SNCA-TG caecum and colon samples (Fig. 2A). Most of the reads (~90%) represented the Firmicutes and Bacteroidetes, which are typically the dominant phyla in the microbiome and belong to the strict anaerobic bacterial group^64^ (Suppl. Fig. 2B and Suppl. Fig. 3B). The phylum Firmicutes was significantly lower in SNCA-TG animals compared with WT in the caecum, but no significant difference was observed in the colon (Suppl. Fig. 1B and Suppl. Fig.2B). In both, the caecum and the colon, SNCA-TG mice had lower levels of Firmicutes/Bacteroidetes ratio (FBR) (Suppl. Fig. 2B and Suppl. Fig. 3B).

**Fig. 2.**
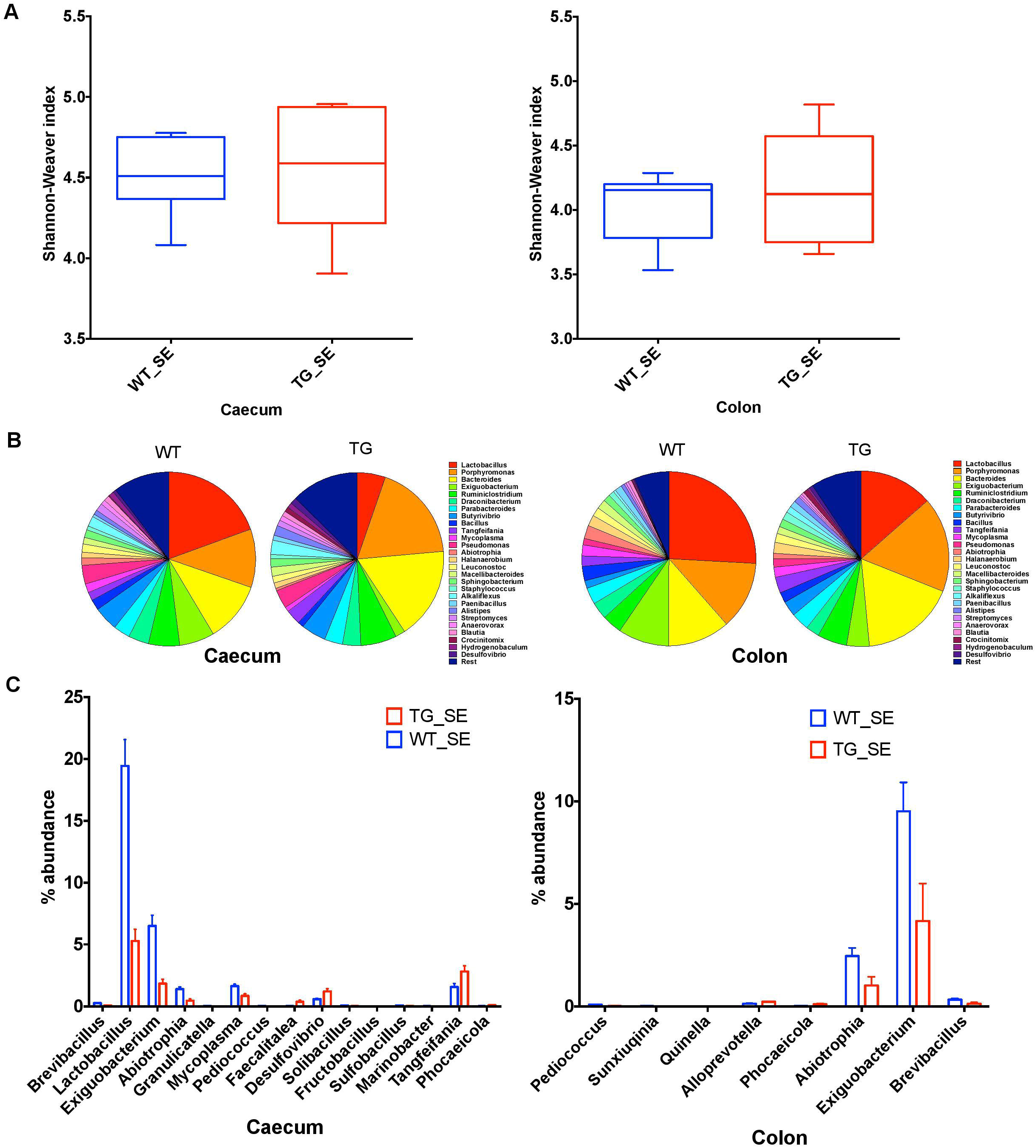
Gut dysbiosis in the caecum and colon of SNCA-TG mice under SE conditions. To identify the gut dysbiosis, 16S rRNA sequencing was performed and α-diversity (Shannon-Weaver index) was measured using MEGAN-CE for the caecum and colon. (A) Shannon-Weaver index was almost similar in WT and SNCA-TG in either caecum or colon. (B) The bacterial abundance at genus level (left figure is for ceacum-WT and TG and right figure is for colon – WT and TG) was calculated in percentage in the caecum and colon. Top 1% bacterial abundance is shown here (32/543) genera. (C) In the caecum 15 different bacterial genera (Brevibacillus, Lactobacillus, Exiguobacterium, Abiotrophia, Granulicatella, Mycoplasma, Pedicoccus, Faecalitalea, Desufovibrio, Solibacillus, Fructobacillus, Sulfobacillus, Marinobacter, Tangfeifania and Phocaeicola) were significantly different between WT and SNCA-TG mice. However, in the colon only 8 genera (Pedicoccus, Sunxiuqinia, Quinella, Alloprevotella, Phocaeicola, Abiotrophia, Exiguobacterium, Brevibacillus) were different in the colon in between WT and SNCA-TG mice. The bacterial genera (Pedicoccus, Phocaeicola, Abiotrophia, Exiguobacterium, Brevibacillus) were common in between the caecum and colon and were significantly different between WT and SNCA-TG mice. Unpaired two-tailed t test was performed the significance measurements for the caecum and colon independently. Data are presented in decreasing p values. P value was considered significance when it was less than or equal to 0.05 (p ≤ 0.05*, ** p≤0.01, *** p≤0.001).

We then investigated differences in taxa among microbial communities (beta-diversity)^65^. We used principal component analysis (PCoA) tool of Megan6^57^ to investigate SNCA-TG and WT samples based on beta diversity metrics which showed a slight shift in clustering for both the groups in the caecum and colon under SE conditions, respectively (Suppl. Fig. 4). We performed an explorative analysis based on the 28 most abundant bacteria genera (more than 1% abundance) and representing more than ~88-92% of the total bacteria sequenced in the caecum and the colon (Fig. 2C). We identified 15 bacterial genera (Brevibacillus, Lactobacillus, Exiguobacterium, Abiotrophia, Granulicatella, Mycoplasma, Pediococcus, Faecalitalea, Desulphovibrio, Solibacillus, Furcotbacillus, Sulfobacillus, Marinobacter, Tangfeifania, Phocaeicola; decreasing order of p values; student’s t-test) in caecum and 8 bacterial genera (Pediococcus, Sunxiuqinia, Quinella, Phocaeicola, Alloprevotella, Abiotrophia, Exiguobacterium, Brevibacillus; decreasing order of p value; student’s t-test) in colon which were differently represented in SNCA-TG mice under SE conditions (Fig. 2D).

Five bacterial genera (Pediococcus, Abiotrophia, Phocaeicola, Exiguobacterium, and Brevibacillus) were significantly changed in both, the caecum and the colon (Fig. 2C) of SNCA-TG compared to WT mice. Lactobacillus genus was the most abundant in the mouse caecum and colon, thus we looked further in detail for the Lactobacillus genus abundance which was significantly reduced in caecal samples of SNCA-TG mice compared with WT (p=0.0001; Student’s t-test), and in a similar way Lactobacillus was also reduced in the colon of the SNCA-TG mice, although it did not reach significance level (p=0.07; Student’s t-test).

### Microbial diversity under EE conditions in SNCA-TG mice

The impact of the environmental conditions on the microbiota was studied using the enriched conditions described previously^55^. Using 16S rRNA, we measured first the bacterial compositions at phyla and genera levels in EE conditions, and found that alpha diversity based on Shannon-Weaver index (SWI) was slightly reduced but did not reach to a significant level in both the caecum and colon (Fig. 3A). At phylum level, there was no change in the FBR (Suppl. Fig. 2B and Suppl. Fig. 3B). The beta diversity, using the PCoA analysis was recruited and no difference was observed between SNCA-TG and WT in EE conditions, respectively (Suppl. Fig. 4). We further performed an explorative analysis based on the 28 most abundant bacteria genera (more than 1% abundance) and representing more than ~88-92% of the total bacteria sequenced in both the caecum and colon (Figure 3B), similar as in the EE conditions. We identified 5 bacterial genera (Tangfeifania, Sphingobacterium, Bacteroites, Barnesiella, Prolixibacter; decreasing order of p value; student’s t-test) in the caecum, and 12 bacterial genera (Roseburia, Anaerotruncus, Anaerosporobacter, Tannerella, Propionibacerium, Tangfeifania, Lachnoclostridium, Holdemanella, Brevibacterium, Fastidiosipila, Quinella, Succinivibrio; decreasing order of p value; student’s t-test) in the colon which were differently represented gut microbiota of SNCA-TG mice in EE conditions (Fig. 3B, C). Out of 17 significantly different abundance genera, only Tangfeifania genus was commonly present in both the caecum and colon. Lactobacillus, the most abundantly occuring genus, was reduced in SNCA-TG mice compared with WT in EE as in the SE conditions, however, it did not reach the significance level (Fig. 3B).

**Fig. 3.**
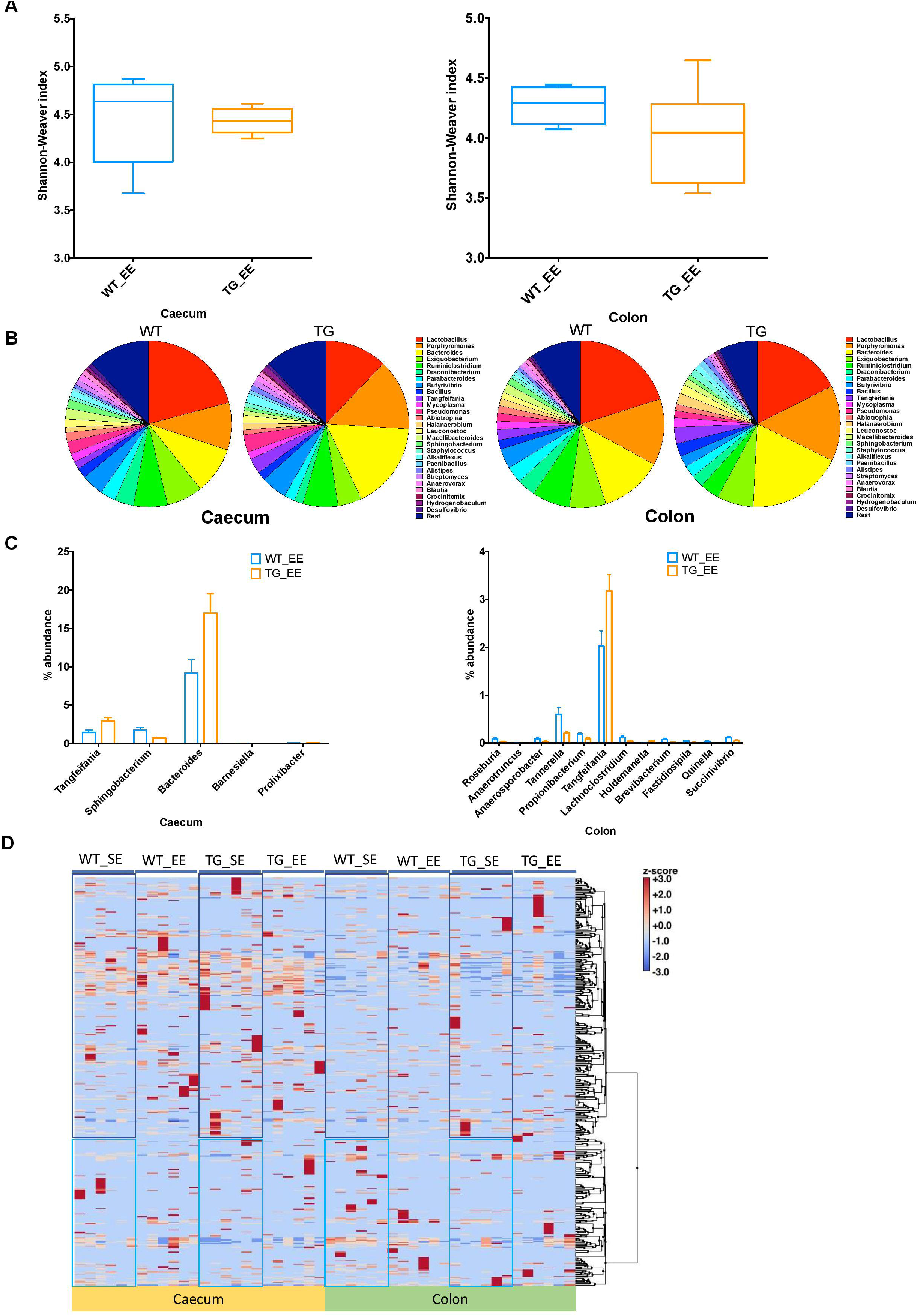
EE conditions moderate the gut dysbiosis in SNCA-TG mice. In a similar fashion to SE conditions, α-diversity (Shannon-Weaver index) was measured in EE conditions using MEGAN-CE for the caecum and colon. (A) Shannon-Weaver index was tending to be lower in SNCA-TG compared with WT in both the caecum and colon. However, it was not significantly different. (B) The bacterial abundance at genus level was calculated in percentage in the caecum (left hand side) and colon (right hand side). (C) EE conditions, levels of the bacterial dysbiosis in the caecum as only 5 different bacterial genera (Tangfeifania, Sphingobacterium, Bacteroides, Barnesiella and Prolixibacter) were significantly different between WT and SNCA-TG mice. However, 12 genera (Roseburia, Anaerotruncus, Anaerosporobacter, Tannerella, Propionibacterium, Tangfeifania, Lachnoclostridium, Holdemanella, Brevibacterium, Fastidiosipila, Quinella and Succinivibrio) were different in the colon for WT and SNCA-TG mice. The only one bacterial genus Tangfeifania was common in between caecum and colon which was significantly different between WT and SNCA-TG mice. Unpaired two-tailed t test was performed the significance measurements for the caecum and colon independently. (D) Overall clustering of bacterial genera is presented in heat map. Abundance of bacterial presence is shown in Z-score (−3.0 to +3.0) values. Two distinct clustering of bacterial genera was appeared in both the caecum and colon in EE and SE conditions. Data are presented in decreasing p values. P value was considered significance when it was less than or equal to 0.05 (p ≤ 0.05*, ** p≤0.01, *** p≤0.001).

### EE conditions affect the bacterial abundance

We calculated the Z-score and clustered bacterial genera for the caecum and colon in the SE and EE conditions (Fig. 4 and Suppl. Fig. 5 & 6). We found that the Z-score for the bacterial abundance was higher in the caecum compared with the colon and that SE and EE conditions have distinct bacterial signature pattern. Further, we measured the alpha diversity using the SWI but found no significant difference between SE and EE conditions (one way ANOVA and Tukey Post-hoc test) in the caecum or colon (Suppl. Fig. 7A). We characterized the most abundant bacterias (28 genera) in the caecum and colon of WT and SNCA-TG mice in both SE and EE environment (Suppl. Fig. 7A & B). Based on the abundance pattern normalized to the environmental conditions, all bacteria were characterized into 6 different groups for the caecum and 5 different groups for the colon (Fig. 4A & B). Furthermore, we standardized percentage bacterial abundance as estimated for SE and EE conditions, and found that EE conditions were able to decrease the percentage gap in the SNCA-TG mice in both the caecum and the colon for the 6 bacterial genera (Fig. 4C). However, in the WT caecum it was almost similar under SE and EE conditions and it was decreased in the colon in the case of EE compared with SE conditions (Fig. 4C).

**Fig. 4.**
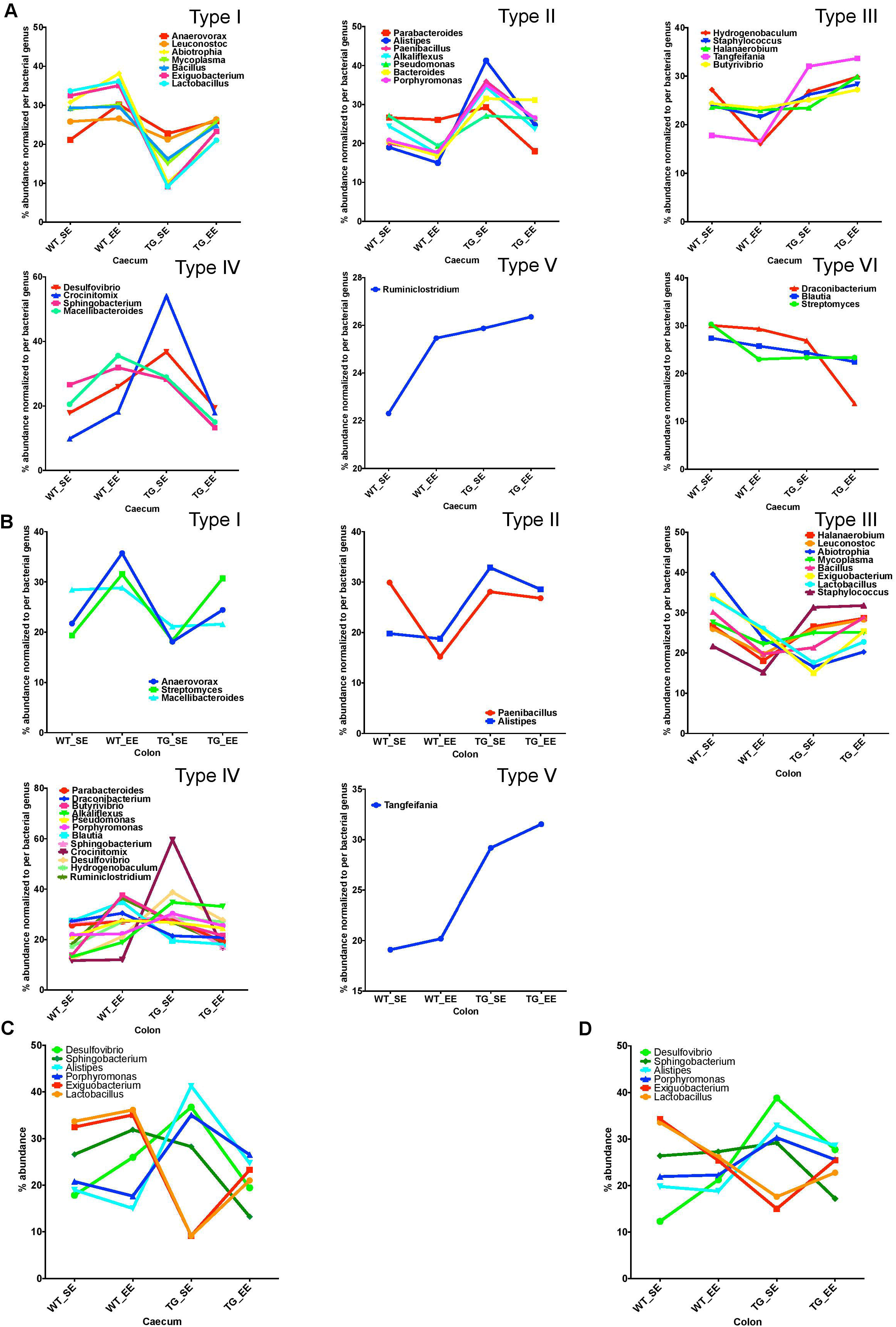
EE conditions promote the growth of anti-inflammatory bacteria. Comparison of SE and EE conditions was done using clustering analysis using MEGEN-CE tool. (A) Bacterial phyla were categorized into 6 different types (Type I-VI) for the caecum. (B) Bacterial phyla were categorized into 6 different types (Type I-VI) for the colon and Type V was missing in the colon for the 32/540 bacterial genera. (C) On the basis of bacterial abundance (normalized for each bacteria for SE and EE conditions), 6 bacterial genera were chosen for the caecum which have two different type (Type I and IV) of abundance. No major change in average abundance was appeared in between WT SE and EE conditions, on the other hand, EE conditions tended to normalize the bacterial abundance in SNCA-TG mice. In the colon, for both the WT and SNCA-TG, EE conditions tended to normalize the bacterial abundance.

### EE condition causes less inflammatory reactions in the colon of SNCA-TG mice

Previous studies in inflammatory bowel disease suggested that inflammatory calprotectin protein is a good marker of inflammation in the gut^66,67^. Calprotectin is secreted by the neutrophils, macrophages and epithelial cells in the gut lumen as a result of inflammation in the colon^67^. Recent studies in PD patients also suggested that calprotectin levels were much higher in patients compared with healthy controls^68^. It appeared that calprotectin could be an inflammatory marker of detection of PD pathology in the feces of the patients^68^ or possibly even in rodent models as described in this study. Therefore, we measured calprotectin expression from digest obtained from the colon from SE and EE conditions in WT and SNCA-TG mice. In SE conditions, SNCA-TG mice had higher levels of calprotectin compared with WT, however, it did not reach the significance level (Fig. 5). On the other hand, in the case of EE conditions (both WT and SNCA-TG mice) calprotectin levels were significantly lower compared to SE conditions. Furthermore, analysis suggested that SNCA-TG mice have significantly less calprotectin in EE compared with SE conditions (Fig. 5), suggesting that EE conditions reduce inflammatory reactions in the gut lumen.

**Fig. 5.**
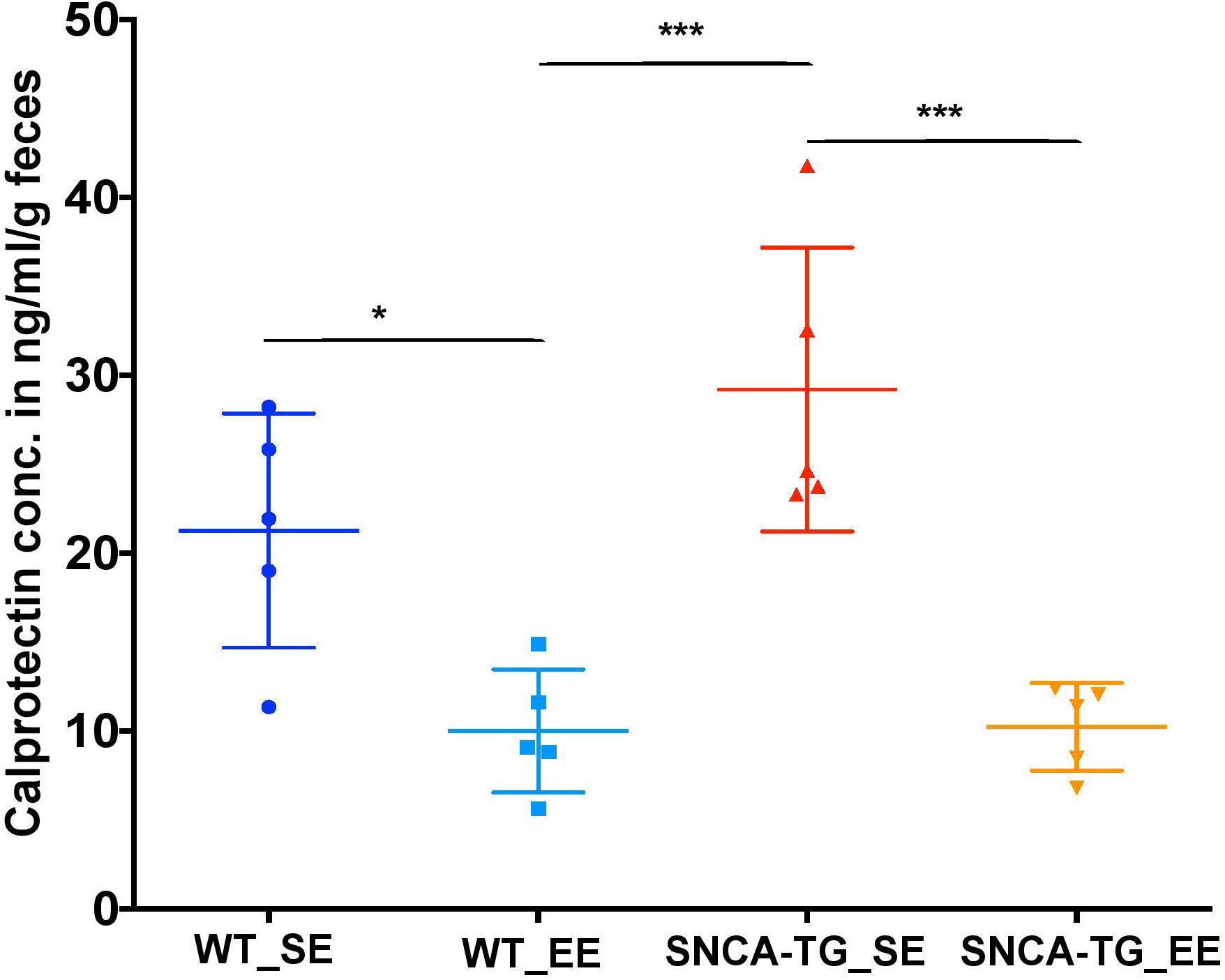
EE conditions dampens the production of inflammatory calprotectin protein. The colon fecal S100A8/S100A9 (Calprotectin, MRP 8/14) protein was estimated using ELISA method. Calprotectin levels were significantly different between SE and EE for the WT and SNCA-TG mice respectively. One-was ANOVA and post-hoc Tukey test was performed to find the significance level between different groups (SE and EE for WT and SNCA-TG). P value was considered significance when it was less than or equal to 0.05 (p ≤ 0.05*, *** p≤0.001).

## Discussion

To study and understand the impact of the gut microbiome in the pathophysiology of PD in patients, complex methodologies using large study groups over long range of time would be required to eliminate factors such as body mass index, age, exposure to antibiotics, diet or ethnicity. Moreover, it would also be critical to find patients in the preclinical stage not experiencing any gastrointestinal or motor symptoms. To tackle these problems, rodent models such as mice or rats are an excellent alternative to identify and perform functional and mechanistic research on host-microbe interactions as they allow manipulation of genome, environment, and gut microbiome composition^69^.

In this study, we are reporting that the microbiome is influenced by the expression of human SNCA in our PD mouse model. Importantly, this study shows that changes in the gut microbiota composition lead to increased inflammatory milieu in the gut in SE conditions in SNCA-Tg animals. Interestingly, a decrease of Lactobacillus observed in our study correlate with a recent report in PD patients and appears to be associated with motor impairment^43,70^. We suggest that these changes may be the consequence of the presence or the accumulation of human α-Syn in the GIT of this animal model rather than the cause of the accumulation. In our study, animals from both genotypes were maintained separately in a conventional standard environment and this change in the gut microbiome abundance could be due to after birth colonization of pups by microbiota (also termed maternal effects). However, multiple studies highlighted that the individuality in gut microbiota composition is more affected by environmental effects such as cohort or litter size than by family^71^. To address this question, we kept the animals in the enriched environmental conditions^55^ and found that indeed enriched conditions are able to affect the gut microbiome composition and inflammation in SNCA-TG mice. Therefore, our studies suggest that the environment plays an important role in the modulation of the microbiome composition in the mouse model despite any potential maternal effects.

Furthermore, it is not really surprising that the microbiota α-diversity in our animal SNCA-TG model was not different from the WT animals as the histological staining did not revealed any acute inflammatory phenotype. It would have been interesting to test, if metabolism, immunity or hormones would have been modulated in the gut of SNCA-TG mice before or after the shift of microbiota to highlight any potential physiological dysregulation of the gut which could have been induced by immunity or metabolism dysfunction. The exact impact of the microbiome in the development of α-synuclein accumulation in the SNCA-TG animals should be now investigated using germ-free conditions or applying broad-spectrum antibiotics. These future studies need to point out whether microbiota can influence the expression levels of α-Syn and if the injection of α-Syn fibrils lead to changes of gut microbial composition similar to the one observed in our animal model.

In summary, we showed that mice overexpressing the complete human *SNCA* gene present an expression of transgenic protein in muscular and mucosal layers of the large intestine. Our study also shows that these animals present a shift of bacterial genera similar to what is observed in PD patients and not related to potential impairment of locomotor activity. All together, we propose that mice overexpressing the human *SNCA* gene represent a good model to study the impact of gut microbiome in connection with the expression of α-Syn.

## Supporting information

EE sensitive gut microbiome in PD_Supp Figs

## Acknowledgements

This is an EU Joint Programme - Neurodegenerative Disease Research (JPND) project. The project JPCOFUND_FP-829-047 aSynProtec is supported through the funding organization Deutschland, Bundesministerium für Bildung und Forschung (BMBF, FKZ) under the aegis of JPND - www.jpnd.eu. This research was funded by the Deutsche Forschungsgemeinschaft (DFG, German Research Foundation) – Projektnummer 407494995 NGS Competent Centre Tübingen (NCCT-DGF). We acknowledge support by Deutsche Forschungsgemeinschaft and Open Access Publishing Fund of University of Tübingen. We would like to acknowledge fruitful discussions with the Institute of Medical Microbiology and Hygiene of Tübingen especially Prof. Julia S Frick and Prof. Matthias Willmann. We also want to thanks the members of the aSynProtec consortium particularly Dr. Jeroen Raes for the help on the detection of calprotectin protein. Funders have no role in study design and data analysis.

## Author’s role

YS: Study design, performed the research, analysed the data, made the figures and wrote the manuscript

MEH, JM: Helped with 16S rRNA data analysis bioinformatics meta data analysis

UK, LQMF: designed, performed and analyzed histology

ZW: performed the animal experiments

JHS: Helped with animal experiments and study design

DH: Helped with study design and data analyses

OR: Study design, funding generation, wrote the manuscript

NC: Study design, performed the research, analysed the data, made the figures and wrote the manuscript

All authors read the manuscript and approved to be co-authors on the manuscript and have substantial contribution in the manuscript.

## Financial Disclosures of all authors (for the preceding 12 months)

This research is supported by JPND grant. NC is supported by NGS Competent Centre grant, YS is supported by JPND consortium grant which was awarded to OR.

---

Suppl. Fig. 1 Human α-Syn expression in GIT tract

Expression of the transgenic human α-Syn protein was investigated by IHC in different parts of the GIT namely stomach, ileum, jejunum, caecum, colon, rectum and found to express the α-Syn protein.

Suppl. Fig. 2 The gut dysbiosis at phylum level in the caecum

To identify the gut dysbiosis, 16S rRNA sequencing was performed to identify (A) the total number of reads (B) % abundance of bacterial phyla (C) and Firmicutes/Bacteroidetes ratio for the caecum.

Suppl. Fig. 3 The gut dysbiosis at phylum level in the colon

To identify the gut dysbiosis, 16S rRNA sequencing was performed to identify (A) the total number of reads (B) % abundance of bacterial phyla (C) and Firmicutes/Bacteroidetes ratio for the colon.

Suppl. Fig. 4 Principal component analysis (PCoA) of the bacterial genera in the caecum and colon

To find the different group similarity or dissimilarity as well as SE and EE condition in the caecum and colon, PCoA was performed. (A) In the caecum, WT and SNCA-TG SE conditions, bacterial genera were differently clustered, however in the case of EE condition, WT and SNCA-TG, clustering of the bacterial genera were less prominent. (B) In the colon, WT and SNCA-TG SE conditions, bacterial genera were overlappingly clustered and similar cluster was also found for the EE conditions.

Suppl. Fig. 5 Clustering of the bacterial genera in the caecum

Comparison of SE and EE conditions was done using clustering analysis using MEGEN-CE tools in the caecum samples.

Suppl. Fig. 6 Clustering of the bacterial genera in the colon

Comparison of SE and EE conditions was done using clustering analysis using MEGEN-CE tools in the colon samples.

Suppl. Fig. 7 Comparison of alpha diversity and abundance of the bacterial phyla in SE and EE conditions

(A) Comparison of the α-diversity for the WT and SNCA-TG in the SE and EE conditions for the caecum and colon samples. (B) The percentage abundance of the bacterial genera for the WT and SNCA-TG in the SE and EE conditions for the caecum and colon samples.

## Notes

Financial Disclosure/Conflict of Interest concerning the research related to the manuscript: None

