## Supplementary figures and images for "Enriched environmental conditions modify the gut microbiome composition and fecal markers of inflammation in Parkinson’s disease"

### EE sensitive gut microbiome in PD_Supp Figs

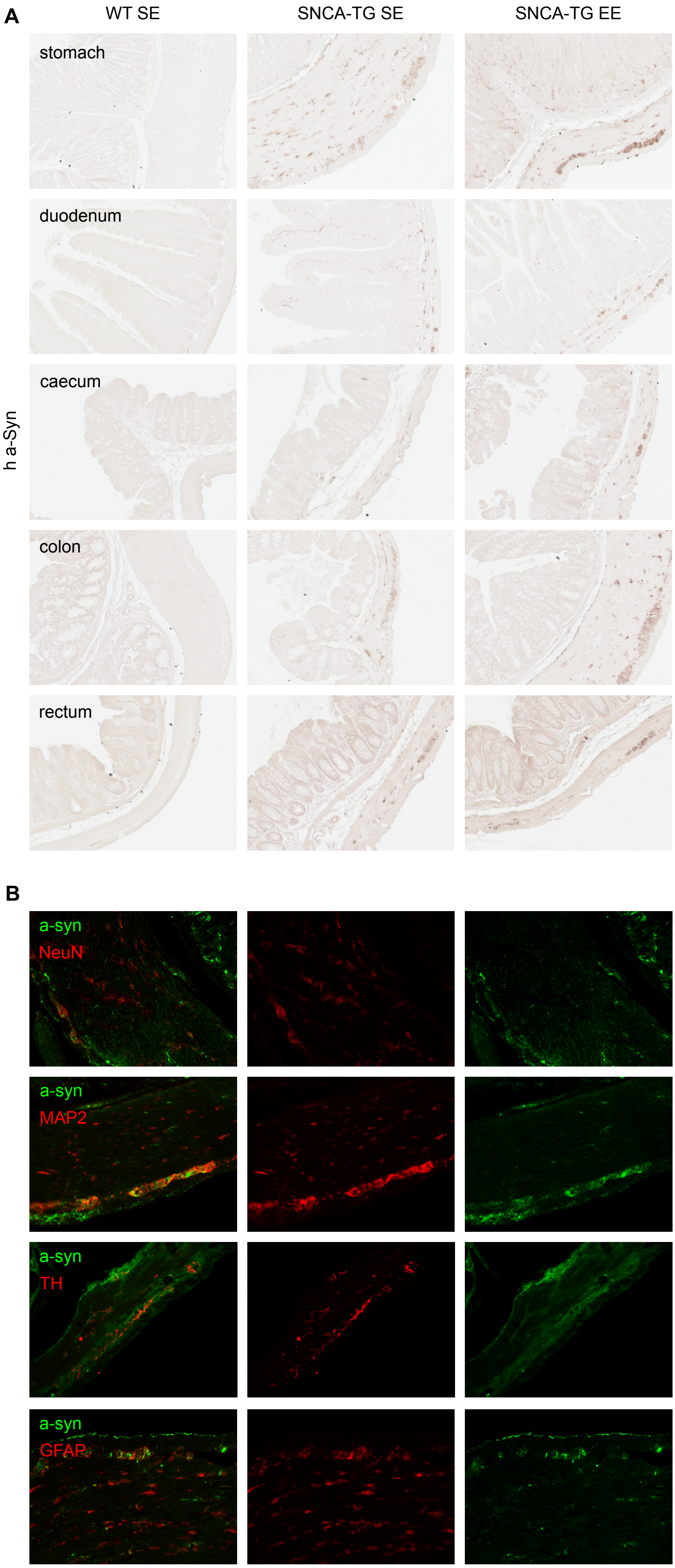

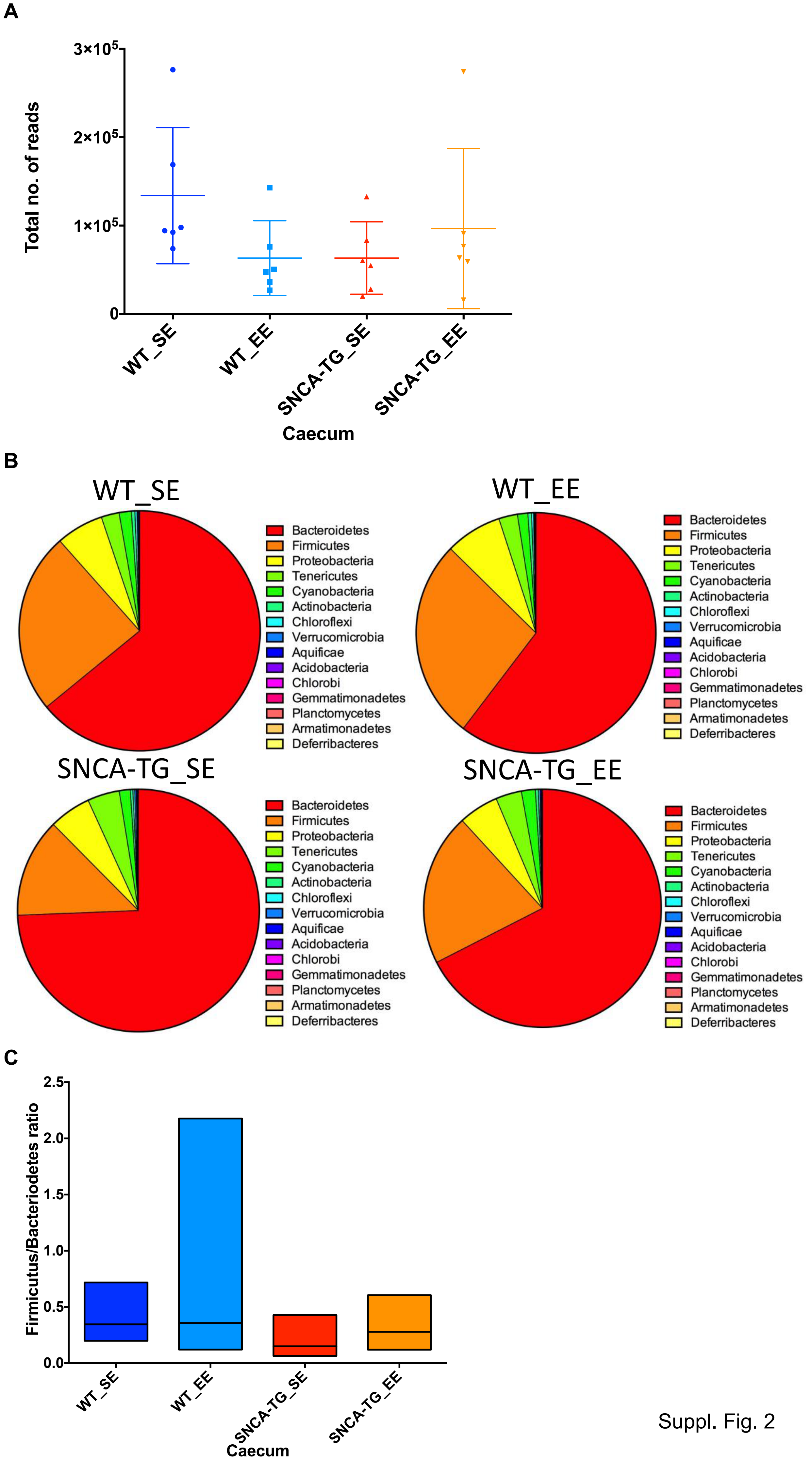

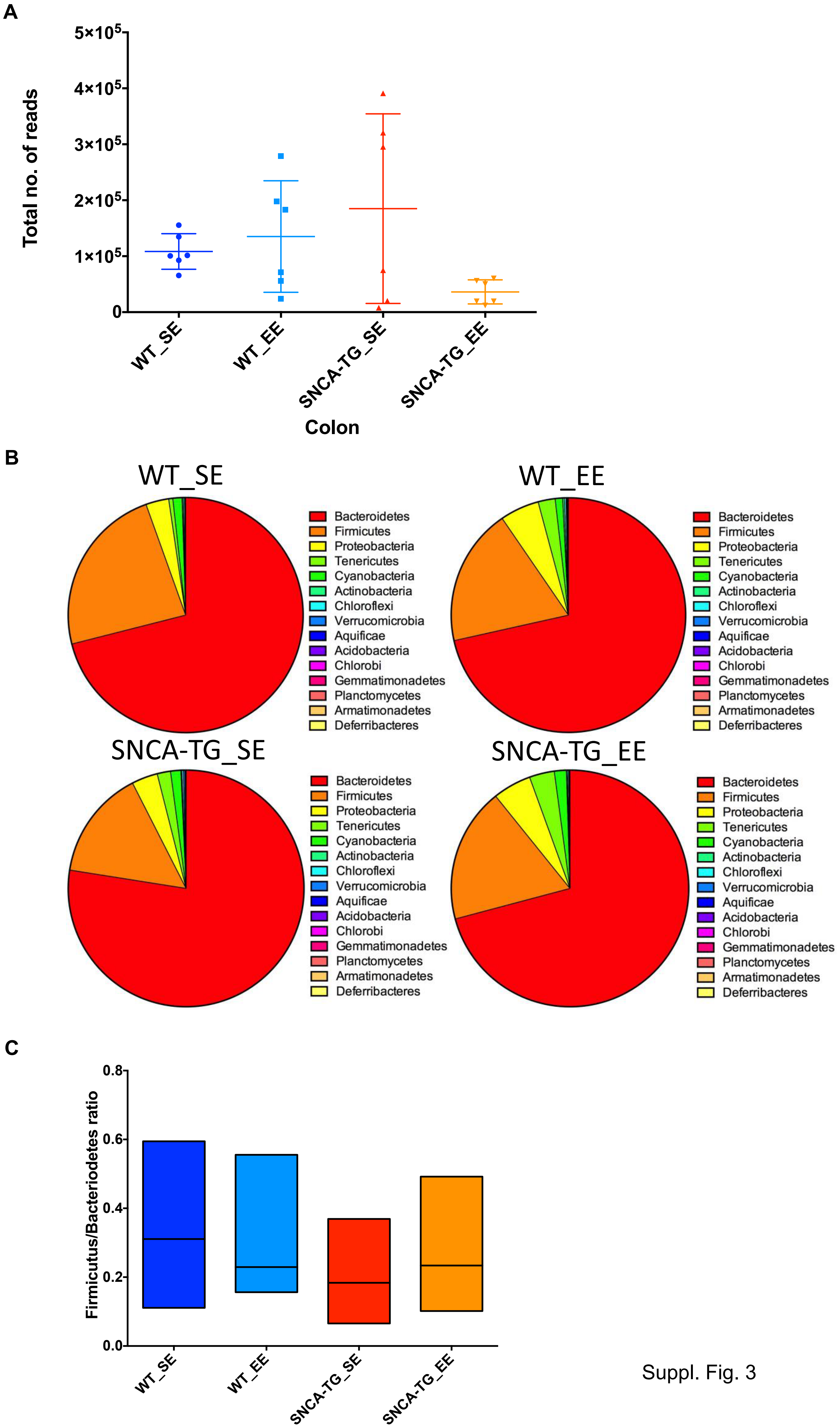

A

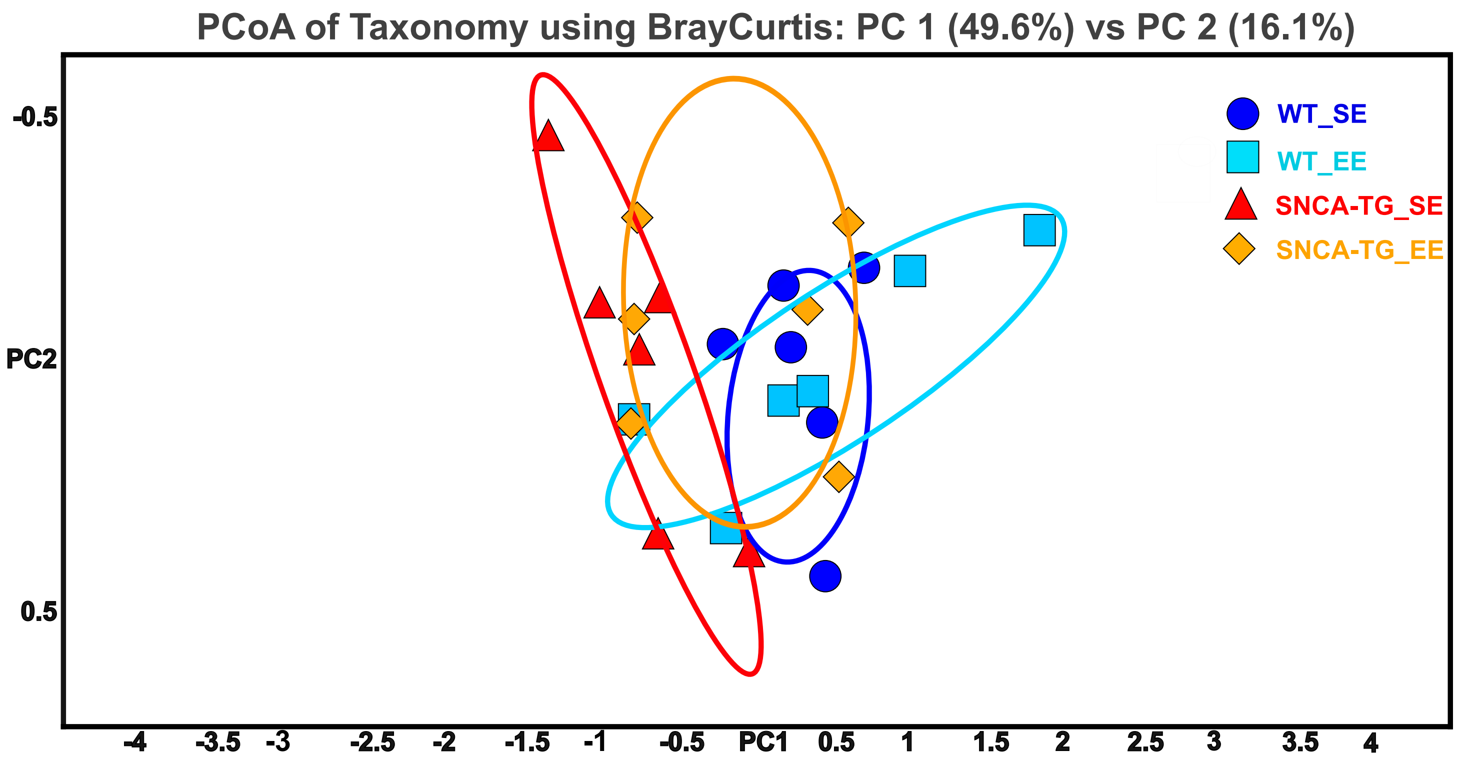


#### Caecum

B

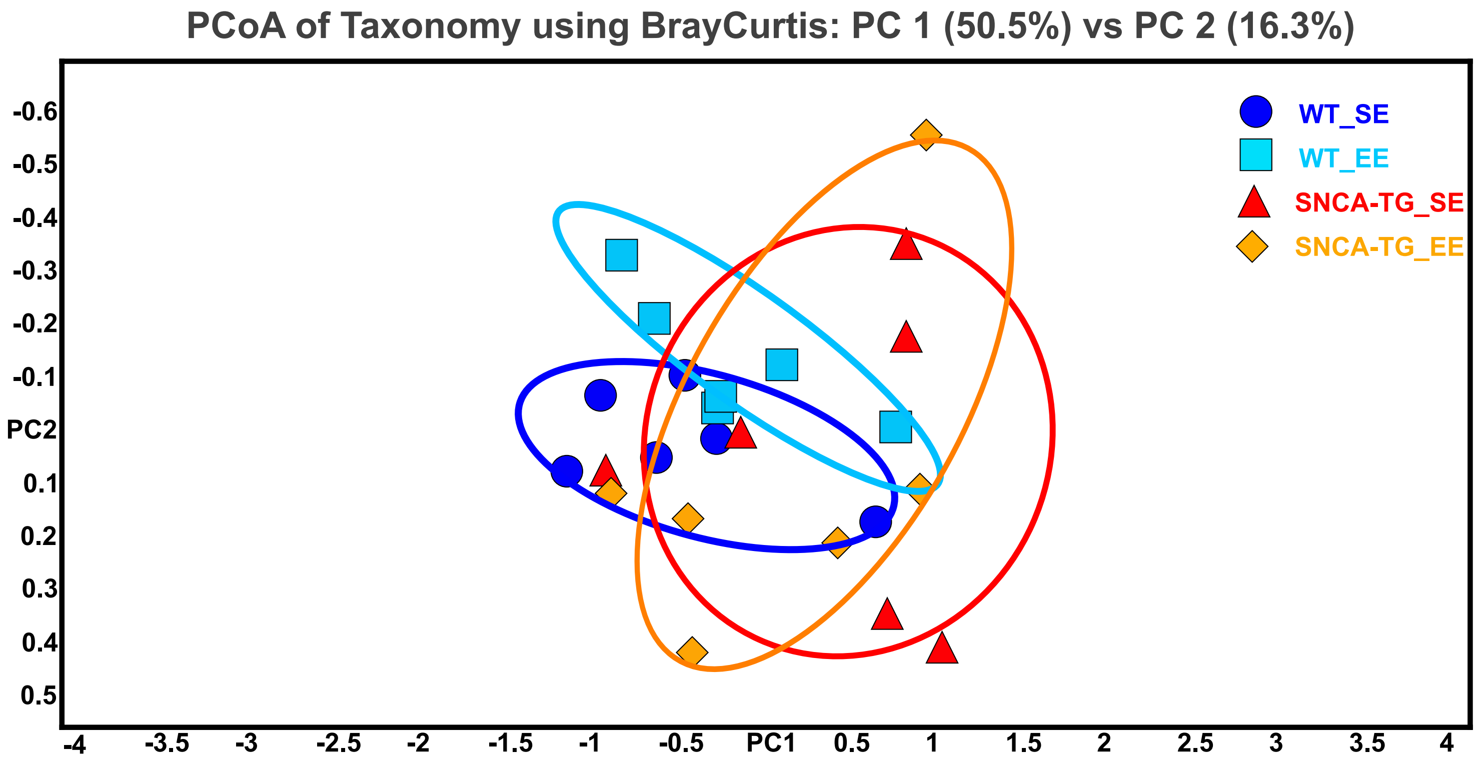


#### Colon

Suppl. Fig. 4

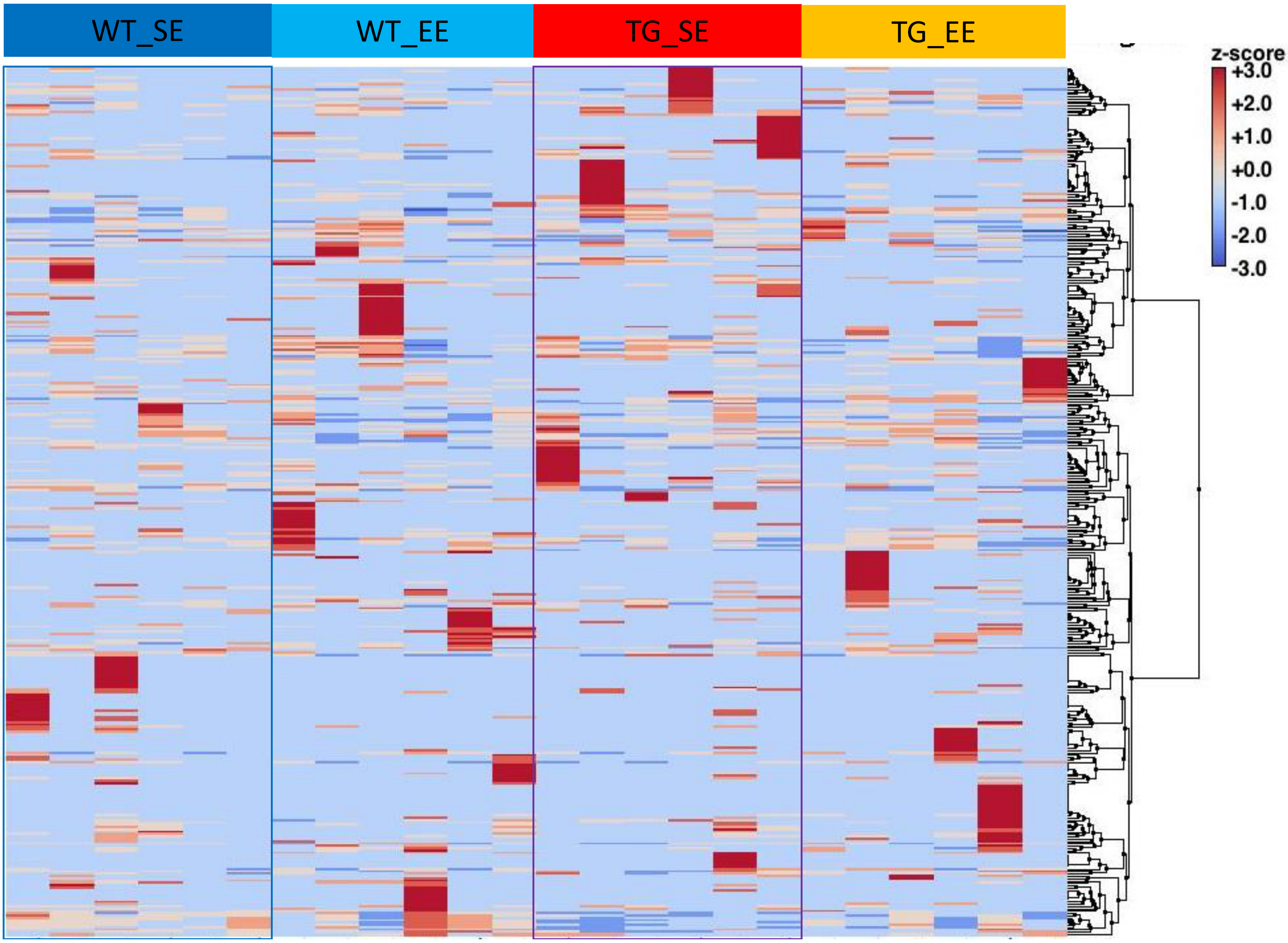


Suppl. Fig. 5

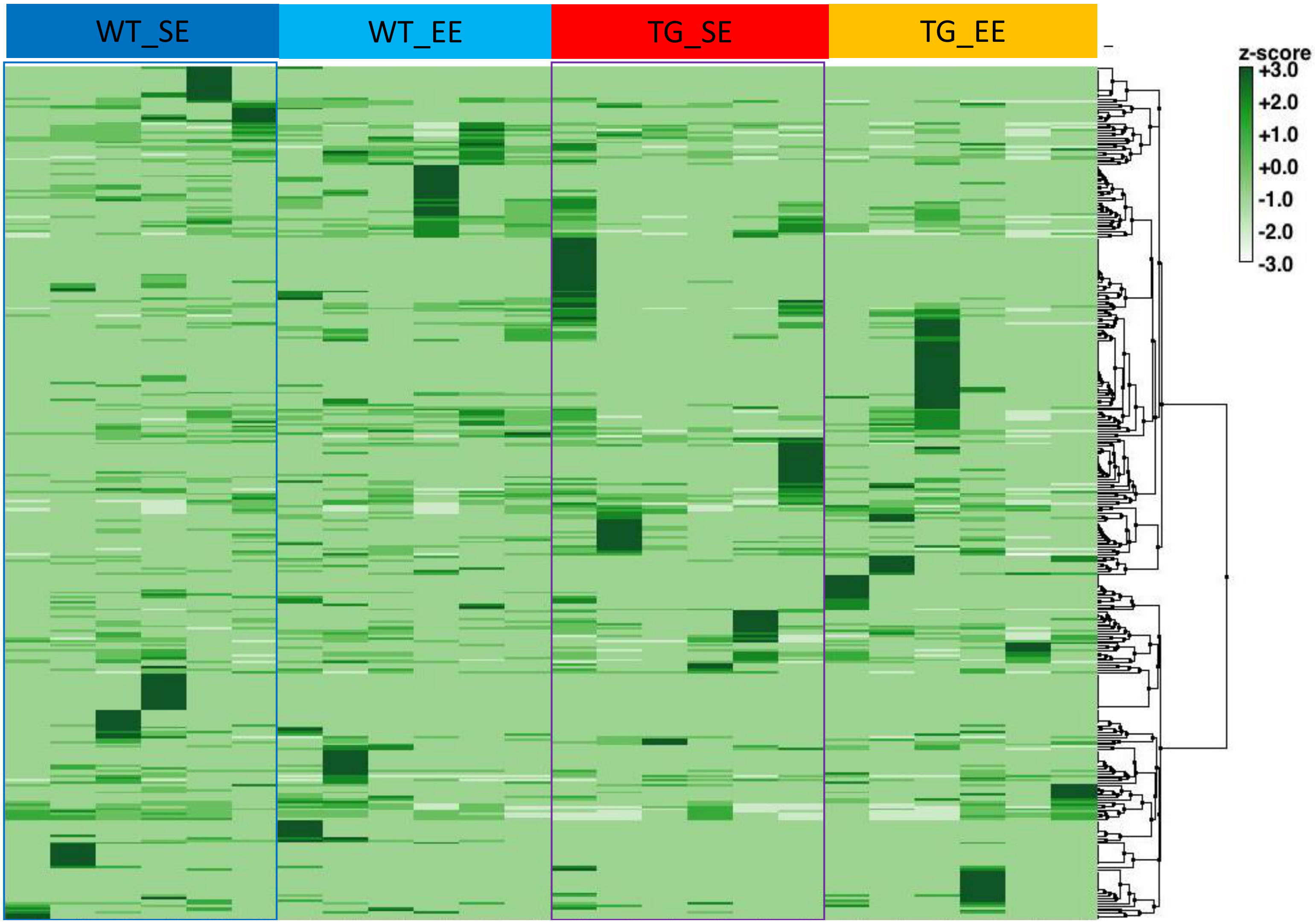


Suppl. Fig. 6

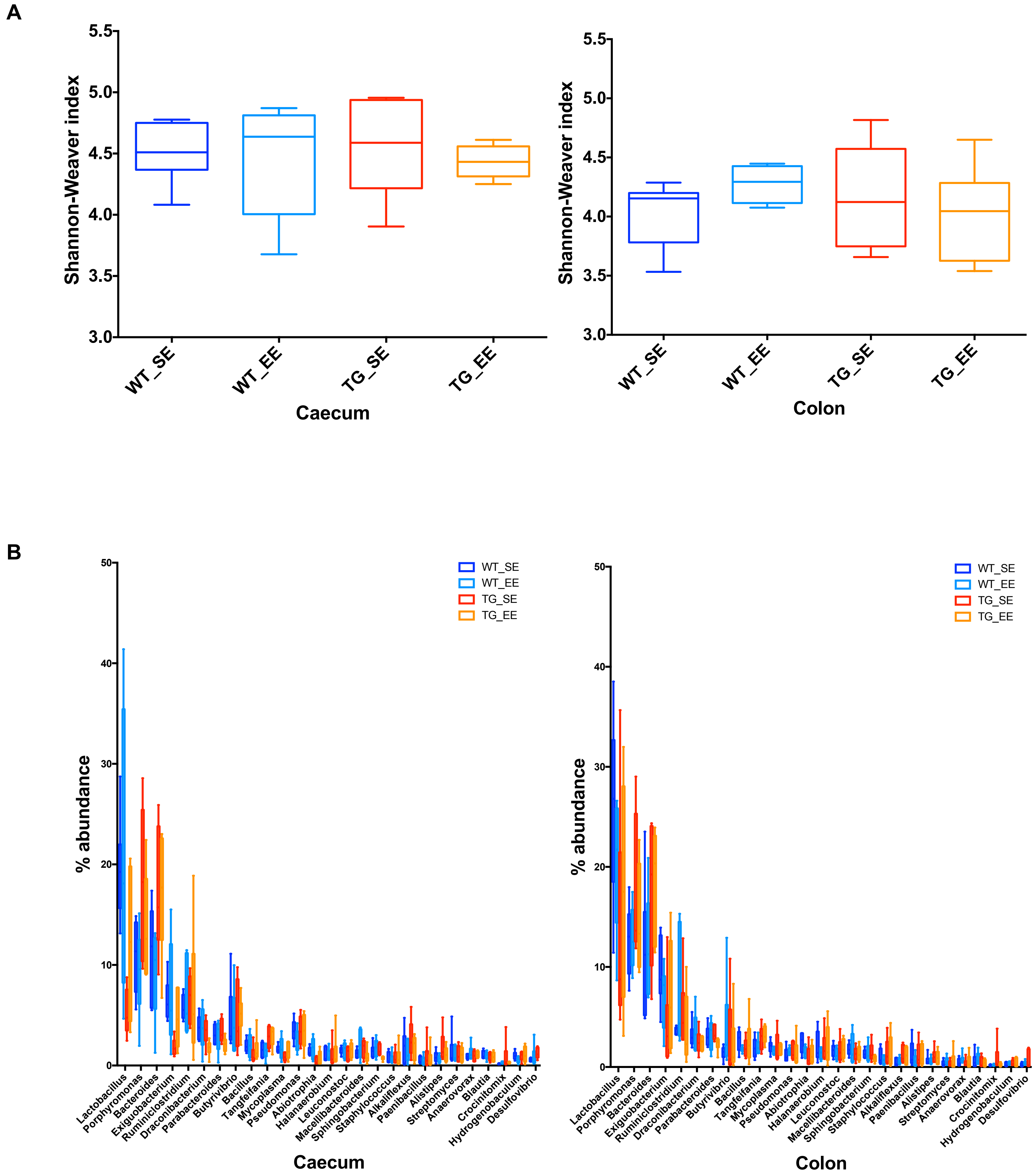
